# Size-selective mortality fosters evolutionary changes in collective learning across ontogeny and decision-making in zebrafish, *Danio rerio*

**DOI:** 10.1101/2022.03.14.484277

**Authors:** Tamal Roy, Tabea Rohr, Robert Arlinghaus

## Abstract

Size-selective harvesting evolutionary alters the life-history, behavioural and physiological traits in exploited fish populations. Changes in these traits may cause alteration in learning and decision-making abilities, either due to energetic trade-offs with brain investment that may vary across development or via correlations with boldness, sociability or other personality traits. Whether size-selective harvesting evolutionarily alters learning and decision-making abilities in fish remains unexplored, despite the global scale of fisheries. We tested the hypothesis that persistent removal of large-bodied individuals typical of many fisheries reduces learning ability in adults but not in juveniles, increases cognitive flexibility but reduces decision-making ability in adults. We examined associative learning through ontogeny, and reversal learning and collective decision-making in adults in three selection lines of zebrafish (*Danio rerio)* generated through positive, negative and random size-selective harvesting for five generations. Fish groups of each selection line were tested across ontogeny using a colour-discrimination paradigm with a food reward. The associative reversal task was conducted with a social reward, and the propensity to make group decisions was tested in an associative task. All selection lines showed significant learning ability and improved performance across ontogeny. Consistent with our hypothesis, the large-harvested line fish revealed a significantly reduced learning speed as subadults and adults, while the small-harvested line fish showed slower error rate compared to controls as 4-month old adults. We found no evidence of memory decay, and the selection lines did not vary in associative reversal ability. Decision-making speed did not vary across lines, but the large-harvested line made faster decisions during the probe trial. We conclude that size-selective harvesting typical of many fisheries evolutionarily alters learning and decision making. As this is likely to negatively and persistently affect resource acquisition and survival in exploited populations, we suggest that the cognition-related mechanism we identify may increase natural mortality.

## Introduction

Intensive harvesting can cause evolutionary changes in animal populations (Allendorf et al. 2009, Heino et al. 2008, Kuparinen et al. 2017). A prominent example of intensive harvesting is fisheries where fish are often targeted based on their body size (Fenberg et al. 2008, Jørgensen et al. 2009, Law 2000). While catching large-sized fish (positive size-selection) is prevalent in most commercial and recreational fisheries, some fishing gears or fisheries governed by certain size-based regulations (e.g., maximum-size limits) may also selectively catch the smaller members of fish populations (negative size-selection) (Heino et al. 2015, Jørgensen et al. 2009, Kuparinen et al. 2009). Both positive and negative size-selection may evolutionarily alter not only the life-history and morphology (Jørgensen et al. 2007, Law 2007, Olsen et al. 2004), but also physiological (Hollins et al. 2018, Redpath et al. 2010, Renneville et al. 2020), and behavioural traits such as boldness (Andersen et al. 2018, Uusi-Heikkilä et al. 2008). What has largely escaped academic attention is that intensive harvesting may also evolutionary affect cognition and learning (Enberg et al. 2012). In this study, we investigate if intensive size-selective mortality fosters evolutionary changes in learning and cognitive decision-making across ontogeny, using zebrafish, *Danio rerio*, as a model for experimental harvesting.

One of the most consistent prediction of theoretical and empirical studies on fisheries-induced evolution is that elevated and positive size-selective harvesting typical of most fisheries fosters the evolution of a fast life-history characterized by early maturation, increased reproductive investment, rapid juvenile, but reduced adult growth, and reduced longevity (Hamilton et al. 2007, Jørgensen et al. 2007, Rijnsdorp 1993). An increased investment in reproduction to fuel a fast life-history can be expected to be traded off with decreased investment in somatic and other energy expensive issues, such as the brain (Energy tradeoff hypothesis: Isler et al. 2009). In turn, increased investment into gonads or reproduction more generally may result in reduced learning, memory and decision-making abilities because investment into brain is positively linked with cognitive ability (Boussard et al. 2021, Buechel et al. 2018, Kotrschal et al. 2015). The consequences for performance might be severe. For example, in guppies *Poecilia reticulata* selected for brain-size, large-brained females outperformed small brained females in a numerical learning task but the large brain lines had smaller guts and produced fewer offspring, suggesting trade-offs with physiological performance (Kotrschal et al. 2013). Increased cognitive abilities also offer fitness benefits with respect to better and quicker access to resources necessary for survival (Dukas 2004). Yet, the relationship between cognition and life-history is not clear because fish demonstrating fast life-histories might also maintain investments into cognition while allocating energy into gonads and reproduction, and achieve this by sacrificing investment into somatic growth and tissue maintenance (Laskowski et al. 2021). Accordingly, fish with fast life-histories may also have larger brains as shown in 21 species of killifish (Sowersby et al. 2021), and brain size might not correspond with improved cognition in all dimensions (Buechel et al. 2018, Fong et al. 2019). It is an open question whether and to what degree size-selective harvesting alters the cognitive ability of evolving fish populations.

In addition to life-history, behavioural types like boldness can also be associated with cognition (Dougherty et al. 2018, Sih et al. 2012). Bolder, more exploratory and active fish are often fast learners and decision makers because they have a higher probability of encountering resources compared to shyer and less active fish (Griffin et al. 2015, Shettleworth 2010). For example in Panamanian bishop fish *Brachyrhaphis episcopi*, individuals that explored more were faster at learning a simple association to access food (DePasquale et al. 2014). In zebrafish, high-predation populations that took higher risks to feed in the presence of predators (Roy et al. 2018b, Roy et al. 2017b) were also faster in learning an associative task and had a longer-lasting memory (Roy et al. 2018a). Size-selective harvesting fosters evolutionary changes in boldness as demonstrated through a range of theoretical (Andersen et al. 2018, Arlinghaus et al. 2017, Claireaux et al. 2018) and empirical (Medaka *Oryzias latipes*: Diaz Pauli et al. 2019, zebrafish: Roy et al. 2021a, Sbragaglia et al. 2020) investigations. Specifically, positive size-selection tends to favour evolution of shy fishes as demonstrated in theoretical (Andersen et al. 2018, Claireaux et al. 2018) as well as empirical studies (Diaz Pauli et al. 2019, Monk et al. 2021). Therefore, it is possible that the cognitive abilities of fish are evolutionarily shifted towards slower learning and reduced memory ability in response to positive size-selection.

Learning abilities may also change throughout ontogeny because of the development of brain with increasing age (Spear et al. 2014). Learning and memory abilities increase with ontogenetic age as has been demonstrated in several fish species (guppies: Boussard et al. 2021, striped knifejaw *Oplegnathus fasciatus*: Makino et al. 2006, jack mackerel *Trachurus japonicus*: Takahashi et al. 2010, zebrafish: Valente et al. 2012). Yet, as investment in brain development has a tradeoff with investment in reproduction (Isler et al. 2009, Kotrschal et al. 2013), shifts in brain development may be associated with the onset of maturation (Buechel et al. 2019). The cognitive abilities change throughout ontogeny because of a transition from resource acquisition fueling growth at the juvenile stage to resource acquisition fueling reproduction when adult (Axelrod et al. 2020, Buechel et al. 2019). Positive size-selection often favours early maturation (Hamilton et al. 2007, Kendall et al. 2014) and this may cause changes in the pace of brain development which may reduce cognitive abilities in adults after the onset of maturation. Further, cognitive abilities are correlated with animal personality (Bensky et al. 2020, Kareklas et al. 2018), and animal personality expression is often consistent within, but not necessarily across life-history stages (Cabrera et al. 2021, Groothuis et al. 2011), as shown in several fish species (killifish *Kryptolebias marmoratus*: Edenbrow et al. 2011, Eastern mosquitofish *Gambusia affinis*: Polverino et al. 2016,zebrafish: Roy et al. 2021a). Hence, a change in personality over ontogeny may be associated with a change in learning ability. Size-selective harvesting causes changes in risk-taking behavior through development in experimentally harvested zebrafish lines (Roy et al. 2021a). In fact, positive size-selection resulted in shy tendencies while negative size-selection resulted in increased boldness in adults (Roy et al. 2021a, Sbragaglia et al. 2020), but not in < 30-day-old juveniles (Roy et al. 2021a). Whether these ontogenetic differences in harvest-induced personality expression are associated with ontogenetic changes in learning ability is currently unexplored and is a major focus of the present work.

Here we tested associative learning ability through ontogeny in three experimental lines of zebrafish generated through positive (large-harvested), negative (small-harvested) and random (control) size-selective harvesting over multiple generations (Uusil□Heikkilä et al. 2015). We predicted that the positive size-selection would result in decreased learning abilities in adults, but not in juveniles, because the large-harvested line showed shy tendencies as adults but not as juveniles (Roy et al. 2021a). We then tested cognitive flexibility among the adults of the selection lines using a reversal learning paradigm. Animals with proactive (bold) coping styles demonstrate less behavioural flexibility than animals with reactive coping (shy) styles (Coppens et al. 2010, Koolhaas et al. 1999). Hence, we expected that the large-harvested line fish that showed shy behavioural tendencies will have increased cognitive flexibility while the small-harvested line that showed increased boldness will have decreased reversal ability (Roy et al. 2021a, Sbragaglia et al. 2020). Finally, we examined possible changes in collective decision-making in adults over time (Hansen et al. 2021, Kareklas et al. 2018) by testing them in an associative task. Fish groups with higher behavioural variability are less cohesive and reach agreement later (Ioannou et al. 2016). As the small- and large-harvested line showed increased and decreased shoal cohesion respectively (Sbragaglia et al. 2021), and decreased and increased variability in boldness (Roy et al. 2021a) compared to the control line, we expected that the small-harvested line will make faster collective decisions while the large-harvested line will make slower collective decisions across repeated trials compared to the controls.

## Materials and methods

### Selection lines

We used three selection lines (large-, small- and random-harvested) of zebrafish, each with a replicate. These lines were generated and maintained as described in Uusi□Heikkilä et al. (2015). The selection lines evolved differences in life-history traits (Uusi□Heikkilä et al. 2015), personality (Roy et al. 2021a, Sbragaglia et al. 2019a, Sbragaglia et al. 2021, Sbragaglia et al. 2020), reproductive behaviour (Roy et al. 2021b, Sbragaglia et al. 2019b), physiological traits (Sbragaglia et al. 2020), and genetics (Sbragaglia et al. 2020, Uusi□Heikkilä et al. 2017). The differences in body-size (Supplementary fig. 1) and other traits were maintained through subsequent generations (F11 to F16) after size-selection was stopped beyond F5 suggesting evolutionary fixation of adaptation to an intensive bout of size-selection (Roy et al. 2021a, Sbragaglia et al. 2021, Sbragaglia et al. 2020).

We used the F_16_ of the three selection lines in the present study, 11 generations after the size-selection had stopped, and fish were reared and maintained without further size-selection. Similar to previous studies (Roy et al. 2021a, Roy et al. 2021b, Sbragaglia et al. 2020), we bred the F_15_ fish of the selection lines in groups (four males and two females), pooled the embryos from each replicate line and stocked eight embryos per line into 30 3-liter boxes (Roy et al. 2021a). We used a total of 240 fish (8 fish × 30 groups; 5 groups per replicate line, 10 groups per selection treatment) for our experiments. The fish were reared and maintained under the following conditions: water temperature 27°C, 12:12h photoperiod; and were fed twice a day with commercial flake food (TetraMin Tropical). We conducted the associative learning assay at four ontogenetic time points (juveniles: 27-38 dpf, subadults: 69-80 dpf, adults: 112-123 and 153-164 dpf) based on Roy et al. (2021a), and reversal and collective decision-making assays in > 6 month old adults (Fig. 1).

**Figure 1:**
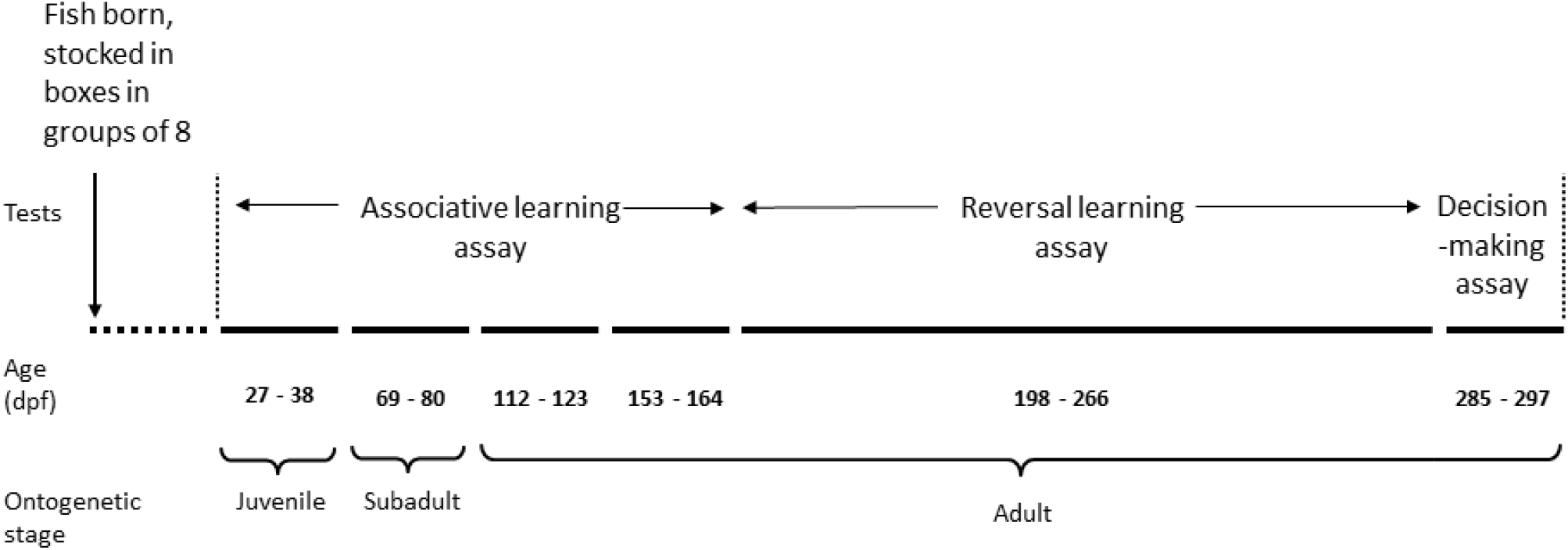
Schematic showing the experimental timeline.

### Associative learning

We tested associative learning and memory ability through ontogeny using a colour discrimination paradigm (Roy et al. 2019, Spence et al. 2011) using 240 fish (8 fish/group, 5 groups/replicate, 10/selection treatment). We used a plus-maze with arm dimensions 27 × 6 × 12 cm, converted it into a T-maze to test juveniles (27-38 dpf) by blocking one of the arms, and increased the length of the arms consecutively by 6 cm for testing subadults (69-80 dpf) and adults (112-123 dpf) (Fig. 2). We tested juveniles, subadults and adults (112-123 dpf) in a two-choice (purple-yellow and blue-brown) discrimination paradigm, and adults between 153-164 dpf in a four-colour discrimination paradigm. The plus maze was used for testing adults at 153-164 dpf age similar to four-choice discrimination paradigms used previously for zebrafish (Roy et al. 2017a, Roy et al. 2018a). Because preferences for primary colours like red and green affect learning abilities in zebrafish (Avdesh et al. 2012, Roy et al. 2019), we did not use these colours unlike previous studies (Roy et al. 2018a, Spence et al. 2011). We trained the fish in groups for six consecutive days and tested their memory on the 12^th^ day without the reward. Removable coloured doors separated the reward chambers from the rest of the maze. In the beginning, we transferred a group of eight fish into the start chamber and allowed them to acclimate for one minute. After this, we released them from the start chamber, and the fish explored the arena for 10 min. We rewarded the fish with food after the first individual entered the correct door. We recorded the trials using an overhead webcam (Logitech B910). After the experiment, we allowed the fish to swim out of the chambers and gently guided them back to the start chamber. We changed the position of the doors randomly during consecutive trials to avoid side bias. From the video recordings, we scored the time taken by a random individual to enter the correct door and commence feeding as the performance time (Hansen et al. 2021, Spence et al. 2011) and the number of incorrect choices (errors) made by the fish before an individual made a correct choice (Kareklas et al. 2018, Roy et al. 2018a).

**Figure 2:**
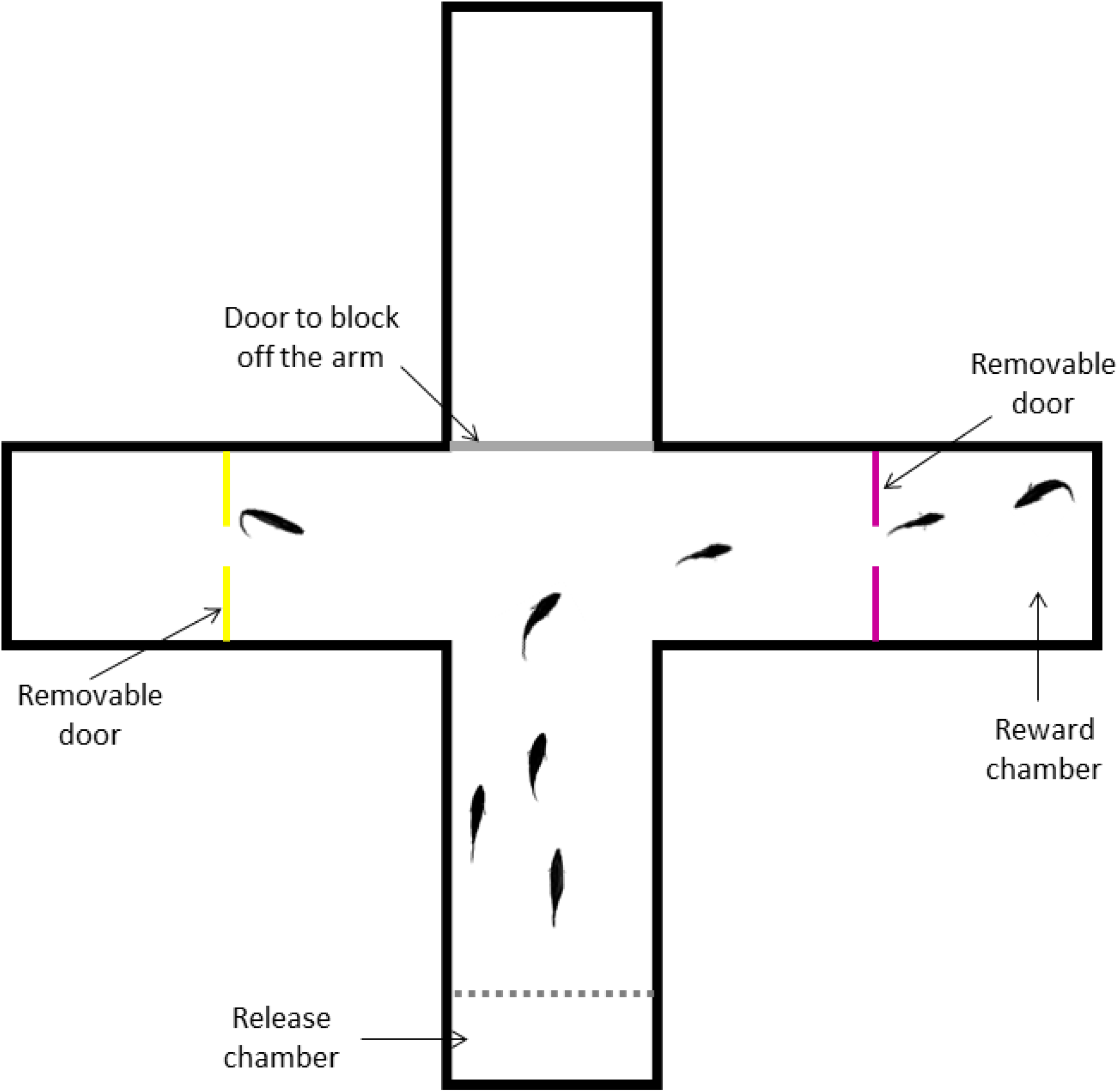
Plus-maze used in associative and reversal learning and collective decision-making assays.

### Reversal learning

We tested reversal learning ability among adult zebrafish (180-190 dpf) in all three selection lines using a purple-yellow discrimination paradigm as described previously, and following the protocol used by Roy et al. (2019). We used fish groups (30 groups, 10 groups/treatment line) and the T-maze from the associative learning assay (Fig. 2), but used a social reward (a group of 10 fish from the same replicate line) instead of a food reward to avoid satiation effects due to repeated testing on the same day (Fernandes et al. 2016, Karnik et al. 2012). We tested half of the groups in purple to yellow reversal and the rest half in yellow to purple reversal. Before the start of the experiments, we transferred 10 individuals (social reward) behind one of the coloured doors and behind a transparent screen (Fig. 2), which allowed the test fish to see them after they swam through the rewarding door. We then moved the test fish group into the start chamber of the T-maze and allowed them to acclimate for one minute. Next, we released the fish from the start chamber and allowed them to explore the arena for 30 minutes. After the trial, we guided the fish back into the start chamber, switched positions of the social reward and the rewarding door and allowed the fish to rest for 10 minutes. We repeated this procedure two more times so that we trained fish in three consecutive 30-minute trials with 10 min rest intervals in between. After the training, we removed the social reward from the arena and changed the side of the rewarding door one last time. We conducted a probe trial for 10 min without the social reward and recorded the trial using an overhead webcam (Logitech B910). We conducted these trials in the morning, allowed the fish to rest in their holding box for 1 h and conducted a second set of training and probe trials in the afternoon by reversing the reward contingency so that there was no reward behind the previously rewarded door and vice versa. From the video recordings, we scored the cumulative time fish spent by fish in the two chambers and estimated the percent cumulative time spent in each chamber (Varga et al. 2018) during the associative (morning) and reversal (afternoon) sessions.

### Collective decision making

We tested collective decision-making among adult zebrafish groups (Kareklas et al. 2018, McAroe et al. 2017) to enter a rewarded (red) door following the same protocol and setup (Fig. 2) as the associative learning assay and using the same fish groups (N=30). We transferred a group of fish into the start chamber and allowed them to acclimate for one min during which we added food to the reward chamber. We released the fish and recorded their behaviour for 5 min. We guided them back to the start chamber after the trial and changed the position of the red door. We trained fish for six consecutive days, and tested for memory on the 13^th^ day without food reward. From the video recordings, we scored the time taken by all fish to enter the rewarded door and commence feeding as the collective decision time (Hansen et al. 2021, Kareklas et al. 2018).

## Statistical analysis

We constructed linear mixed-effects regression models (lmer) to test for change in associative learning ability through ontogeny among selection lines. To test for changes in performance across successive trials during training at different ontogenetic stages, we first transformed the response variable (performance time) using Tukey’s ladders of powers transformation and confirmed the normality and heterogeneity of residuals. We then fitted mixed effects models using the transformed measure as the dependent variable, interaction of ‘Selection line’, ‘Age’ (ontogenetic stage) and ‘Trial’ as the fixed effect and ‘Group ID’ nested within ‘Replicate’ as the random effect and ‘Age’ as random slope. To test for changes in errors made across successive trials over ontogeny, we fitted a generalized linear mixed-effects model (glmer) with a Poisson structure using ‘Errors’ as the dependent variable, interaction of ‘Selection line’, ‘Age’ and ‘Trial’ as the fixed effect, ‘Group ID’ nested within ‘Replicate’ as the random effect and ‘Age’ as the random slope. To test for memory among the selection lines, we compared the performance time and the no. of errors made on the 6^th^ day (last day of training) and the 12^th^ day (probe trial) using paired t-test and Wilcoxon paired-sample test.

For testing differences in reversal learning ability among selection lines, we logit transformed the response variable ‘percent cumulative time’ and compared the transformed percent time spent by fish in correct (rewarded) and incorrect (unrewarded) chambers during associative and reversal trials. A significantly higher percent time spent in the correct than incorrect chamber within the trials indicated associative/reversal learning.

We tested change in time to make correct decisions over consecutive trials among selection lines, and collective memory using linear mixed effects models. We Tukey-transformed ‘collective decision time’ and confirmed the normality and heterogeneity of residuals. We used the transformed measure as the dependent variable, interaction of ‘Selection line’ and ‘Trial’ as the fixed effect and ‘Group ID’ nested within ‘Replicate’ as the random effect. To test for collective memory, we constructed a similar mixed model and compared decision time on 6^th^ and 13^th^ day trials.

All analyses were done in R studio version 3.6.1 (R Development Core Team 2019). Data were transformed using *rcompanion* (Mangiafico et al. 2017) and ‘*car*’ (Fox et al. 2007) packages, and ‘lmer’ and ‘glmer’ models were constructed using *lmerTest* (Kuznetsova et al. 2017) and *lme4* (Bates et al. 2012) packages. Line and box-whisker plots were made using *ggplot2* (Wickham 2011), *plyr* (Wickham et al. 2020) and *ggpubr* (Kassambara et al. 2020) packages in R.

## Results

We found significant differences in associative learning and memory among selection lines and over ontogenetic age. Performance time changed significantly across trials (F_1,614_=27.21, p<0.01) during training, and over ontogenetic age (F_3,15_=10.54, p<0.01) for all selection lines (Table 1a, Fig. 3a). Fish performed significantly better as adults at 112-123 dpf (t=-2.03, p=0.08) and 154-164 dpf (t=-3.21, p<0.01) age compared to juveniles at 27-38 dpf age (Table 1b). Errors made by fish decreased significantly with training (z=-5.52, p<0.01), and adults at 112-123 dpf made significantly less errors (z=-7.00, p<0.01) across trials compared to juveniles (Table 2, Fig. 3b). The large-harvested line fish took significantly more time to find the reward over successive trials as subadults at 69-80 dpf (t=2.12, p=0.03), and adults at 112-123 dpf (t=2.65, p<0.01), compared to the control line (Fig. 3a). The performance time over successive trials of the small-harvested line fish did not differ significantly from the controls at any ontogenetic time-point (Table 1b, Fig. 3a). Errors made across trials were significantly higher in subadults at 69-80 dpf (z=2.21, p=0.03) and adults at 112-123 dpf (z=9.31, p<0.01) in the large-harvested line compared to controls. Errors made across trials were significantly higher as adults at 112-123 dpf (z=2.35, p=0.02) but significantly lower at 154-164 dpf (z=-3.78, p<0.01) of the small-harvested line compared to controls (Table 2, Fig. 3b). These results meant that the subadults and adults of the large-harvested line showed a significantly slower improvement in performance during training (Fig. 3a, b), and the small-harvested line made more errors as 4-month old adults, compared to the control line (Fig. 3a, b). In tests for memory, we found no significant differences in performance time (Table 3a, Fig. 4a) and errors made (Table 3b, Fig. 4b) between the 6^th^ and the probe (12^th^ day) trials, indicating similar retention of memory in all selection lines (Table 3a, b, Fig. 4a b).

**Table 1.** Results of (a) main effects and (b) fixed effects obtained from lmer model comparing performance time across trials of LH (large-harvested) and SH (small-harvested) lines with the control line across ontogenetic stages A (juvenile) to D (adult). Significant results are in bold (marginal: ‘^+^’).

| (a) Main effects | Sum Sq | Mean Sq | NumDF | DenDF | F-value | Pr(>F) |
| --- | --- | --- | --- | --- | --- | --- |
| Selection line | 0.002 | 0.001 | 2 | 4.06 | 0.68 | 0.56 |
| Trial | 0.037 | 0.037 | 1 | 614.06 | 27.21 | **< 0.001** |
| Age | 0.043 | 0.014 | 3 | 14.98 | 10.54 | **< 0.001** |
| Selection line × Trial | 0.006 | 0.003 | 2 | 614.06 | 2.21 | 0.11 |
| Selection line × Age | 0.007 | 0.001 | 6 | 14.98 | 0.87 | 0.54 |
| Trial × Age | 0.001 | 0.000 | 3 | 614.06 | 0.25 | 0.86 |
| Selection line × Trial × Age | 0.015 | 0.002 | 6 | 614.06 | 1.78 | 0.10 |

| (b) Fixed effects | Estimate | SE | df | t-value | Pr(>\|t\|) |
| --- | --- | --- | --- | --- | --- |
| Intercept | -6.61e-01 | 1.30e-02 | 2.81e+01 | -50.66 | **< 0.001** |
| Selection line: LH | -5.98e-03 | 1.85e-02 | 2.81e+01 | -0.32 | 0.75 |
| Selection line: SH | -1.02e-02 | 1.85e-02 | 2.81e+01 | -0.55 | 0.59 |
| Trial | -1.50e-03 | 2.80e-03 | 6.14e+02 | -0.54 | 0.59 |
| Stage B (69-80 dpf) | -4.99e-03 | 2.08e-02 | 1.09e+01 | -0.24 | 0.81 |
| Stage C (112-123 dpf) | -4.99e-02 | 2.45e-02 | 6.95e+00 | -2.03 | 0.08<sup>+</sup> |
| Stage D (153-164 dpf) | -6.83e-02 | 2.12e-02 | 9.82e+00 | -3.21 | **< 0.01** |
| LH × Trial | -4.15e-03 | 3.95e-03 | 6.14e+02 | -1.05 | 0.29 |
| SH × Trial | -2.19e-03 | 3.95e-03 | 6.14e+02 | -0.55 | 0.58 |
| LH × Stage B | 2.37e-02 | 2.94e-02 | 1.09e+01 | -0.81 | 0.44 |
| SH × Stage B | 2.37e-02 | 2.94e-02 | 1.09e+01 | -0.94 | 0.37 |
| LH × Stage C | -1.64e-02 | 3.47e-02 | 6.95e+00 | -0.47 | 0.65 |
| SH × Stage C | -2.90e-02 | 3.47e-02 | 6.95e+00 | -0.84 | 0.43 |
| LH × Stage D | 1.85e-02 | 3.00e-02 | 9.82e+00 | 0.62 | 0.55 |
| SH × Stage D | 1.07e-02 | 3.00e-02 | 9.82e+00 | 0.36 | 0.73 |
| Trial × Stage B | -7.40e-03 | 3.95e-03 | 6.14e+02 | -1.87 | 0.06<sup>+</sup> |
| Trial × Stage C | -6.65e-03 | 3.95e-03 | 6.14e+02 | -1.68 | 0.09<sup>+</sup> |
| Trial × Stage D | -7.88e-04 | 3.95e-03 | 6.14e+02 | -0.20 | 0.84 |
| LH × Trial × Stage B | 1.19e-02 | 5.59e-03 | 6.14e+02 | 2.12 | **0.03** |
| SH × Trial × Stage B | 5.38e-03 | 5.59e-03 | 6.14e+02 | 0.96 | 0.34 |
| LH × Trial × Stage C | 1.48e-02 | 5.59e-03 | 6.14e+02 | 2.65 | **<0.01** |
| SH × Trial × Stage C | 5.31e-03 | 5.59e-03 | 6.14e+02 | 0.95 | 0.34 |
| LH × Trial × Stage D | 3.50e-03 | 5.59e-03 | 6.14e+02 | 0.63 | 0.53 |
| SH × Trial × Stage D | -3.46e-03 | 5.59e-03 | -6.14e+02 | -0.62 | 0.54 |

**Table 2.** Results of fixed effects from glmer model comparing the number of mistakes made across trials by large-and small-harvested line, with the control line across stages A (juvenile) to D (adult). Significant results are in bold (marginal: ‘^+^’).

| Fixed effects | Estimate | SE | z-value | Pr(>\|t\|) |
| --- | --- | --- | --- | --- |
| Intercept | 1.66 | 0.30 | 5.45 | < **0.001** |
| Selection line: LH | -0.04 | 0.43 | -0.09 | 0.93 |
| Selection line: SH | -1.09 | 0.45 | -2.41 | **0.02** |
| Trial | -0.23 | 0.04 | -5.52 | < **0.001** |
| Stage B (69-80 dpf) | 0.93 | 0.36 | 2.58 | **0.01** |
| Stage C (112-123 dpf) | 0.91 | 0.58 | 1.58 | 0.11 |
| Stage D (153-164 dpf) | -0.81 | 0.44 | -1.83 | 0.07<sup>+</sup> |
| LH × Trial | -0.07 | 0.07 | -1.11 | 0.27 |
| SH × Trial | 0.19 | 0.07 | 2.87 | < **0.01** |
| LH × Stage B | -0.40 | 0.51 | -0.77 | 0.44 |
| SH × Stage B | 0.41 | 0.53 | 0.77 | 0.44 |
| LH × Stage C | -2.95 | 0.84 | -3.51 | < **0.001** |
| SH × Stage C | 0.42 | 0.83 | 0.51 | 0.61 |
| LH × Stage D | 0.31 | 0.63 | 0.49 | 0.62 |
| SH × Stage D | 2.10 | 0.63 | 3.32 | < **0.001** |
| Trial × Stage B | -0.03 | 0.05 | -0.60 | 0.55 |
| Trial × Stage C | -0.56 | 0.08 | -6.98 | < **0.001** |
| Trial × Stage D | 0.28 | 0.05 | 5.01 | < **0.001** |
| LH × Trial × Stage B | 0.17 | 0.08 | 2.21 | **0.03** |
| SH × Trial × Stage B | -0.01 | 0.08 | -0.11 | 0.91 |
| LH × Trial × Stage C | 1.01 | 0.11 | 9.31 | < **0.001** |
| SH × Trial × Stage C | 0.25 | 0.11 | 2.35 | **0.02** |
| LH × Trial × Stage D | -0.13 | 0.09 | -1.43 | 0.15 |
| SH × Trial × Stage D | -0.32 | 0.08 | -3.78 | < **0.001** |

**Table 3.**
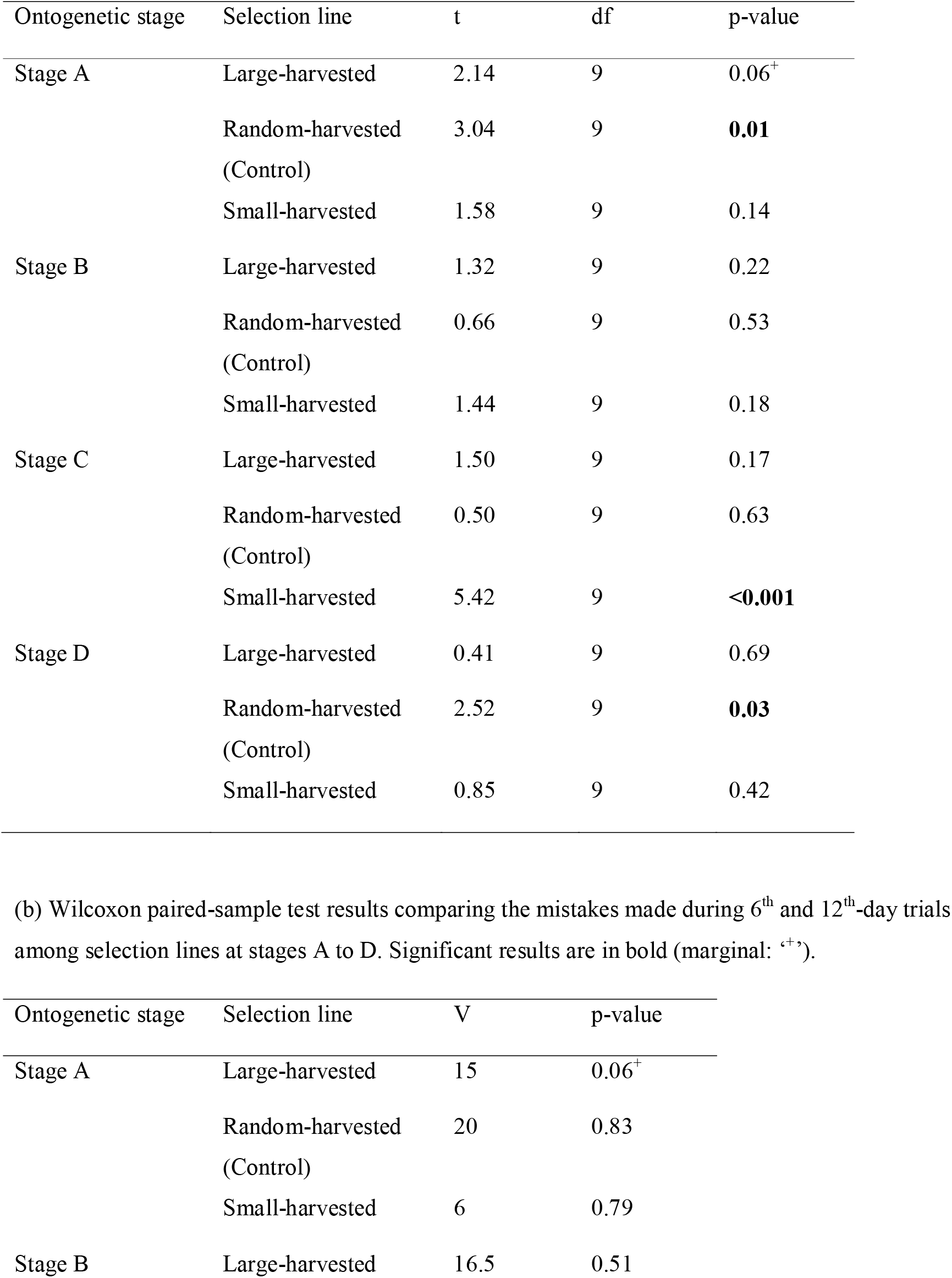
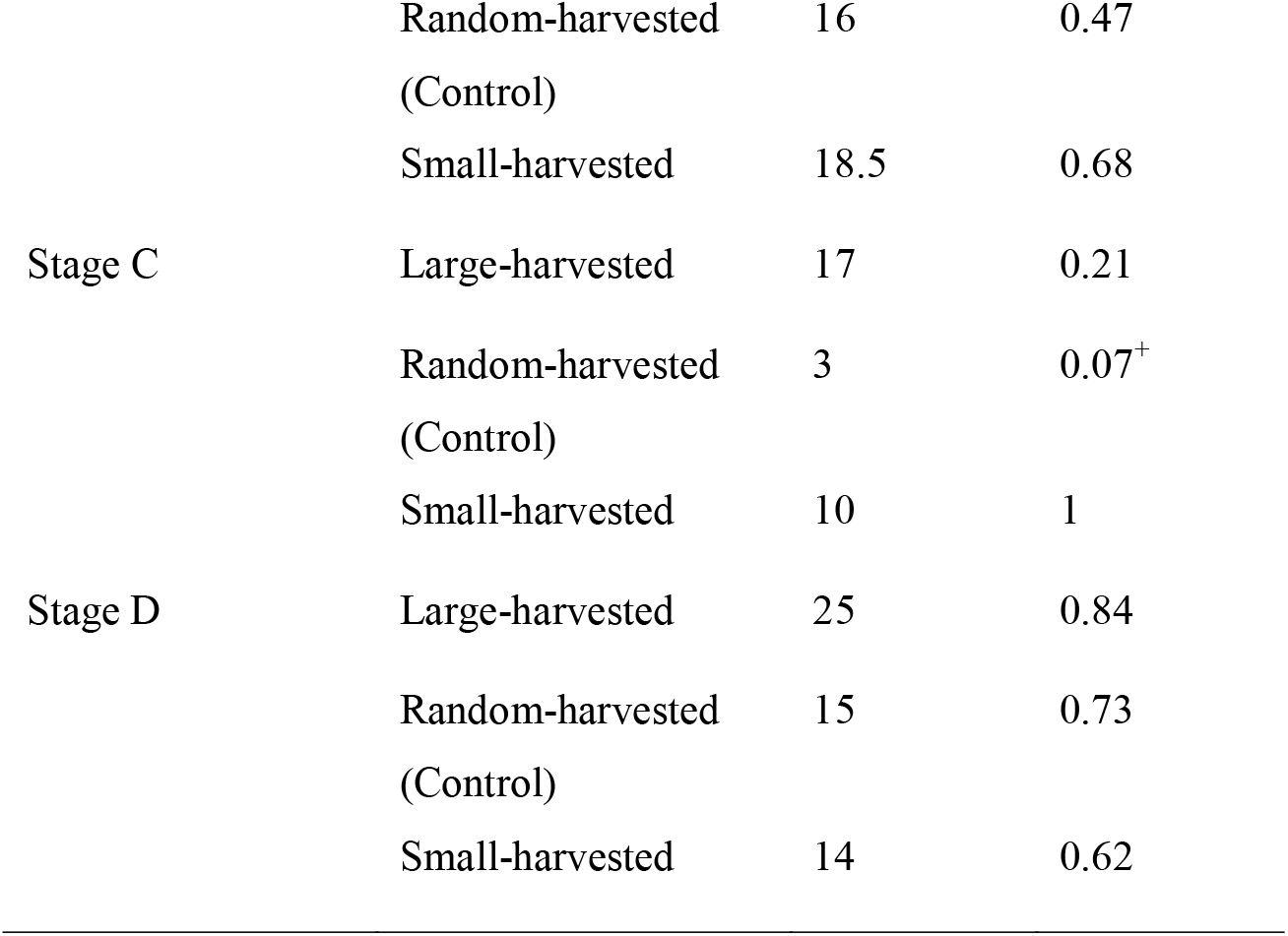
Evaluating memory performance. (a) Paired t-test results comparing performance time of fish during 6^th^ and 12^th^-day trials among selection lines at four ontogenetic stages A (27-38 dpf), B (69-80 dpf), C (112-123 dpf) and D (153-164 dpf). Significant results are in bold (marginal: ‘^+^’).(b) Wilcoxon paired-sample test results comparing the mistakes made during 6^th^ and 12^th^-day trials among selection lines at stages A to D.Significant results are in bold (marginal: ‘^+^’)

**Figure 3a:**
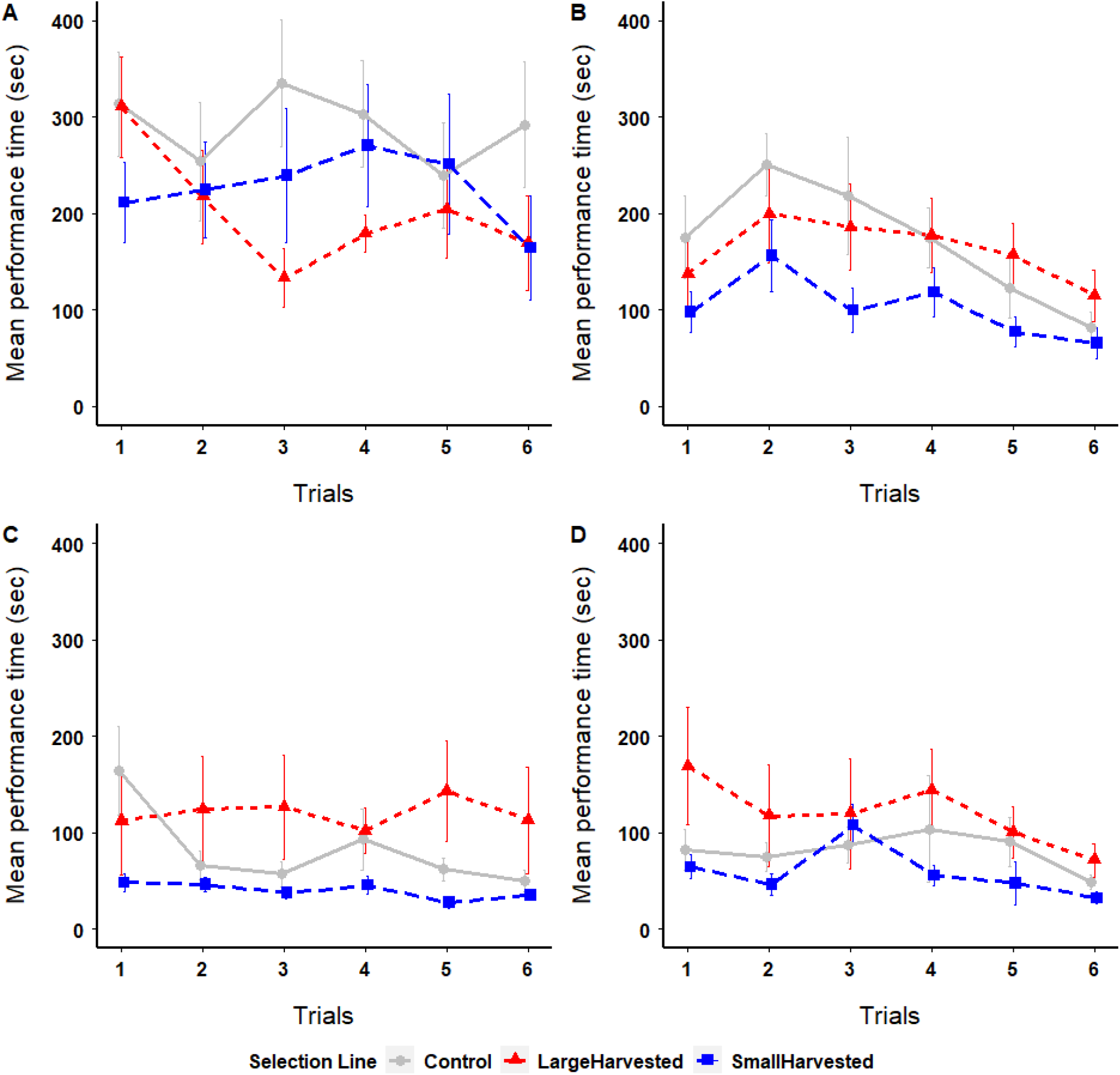
Comparison of mean (±SE) performance time (sec) during training as (**A**) juveniles (27-38 dpf), (**B**) subadults (69-80 dpf) and adults at (**C**) 112-123 dpf, and (**D**) 153-164 dpf

**Figure 3b:**
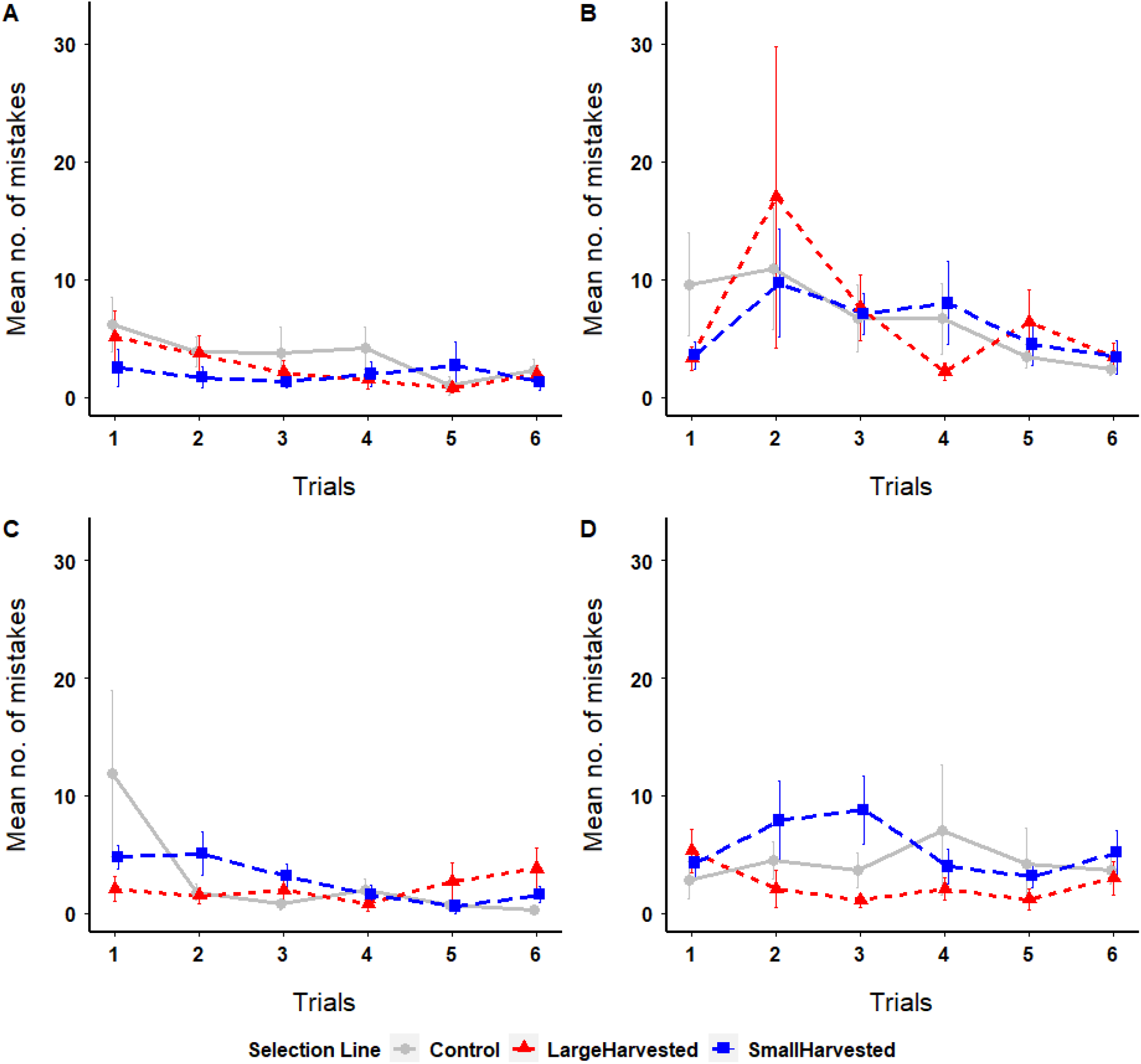
Comparison of mean number of mistakes made across trials as (**A**) juveniles (27-38 dpf), (**B**) subadults (69-80 dpf) and adults at (**C**) 112-123 dpf, and (**D**) 153-164 dpf

**Figure 4a:**
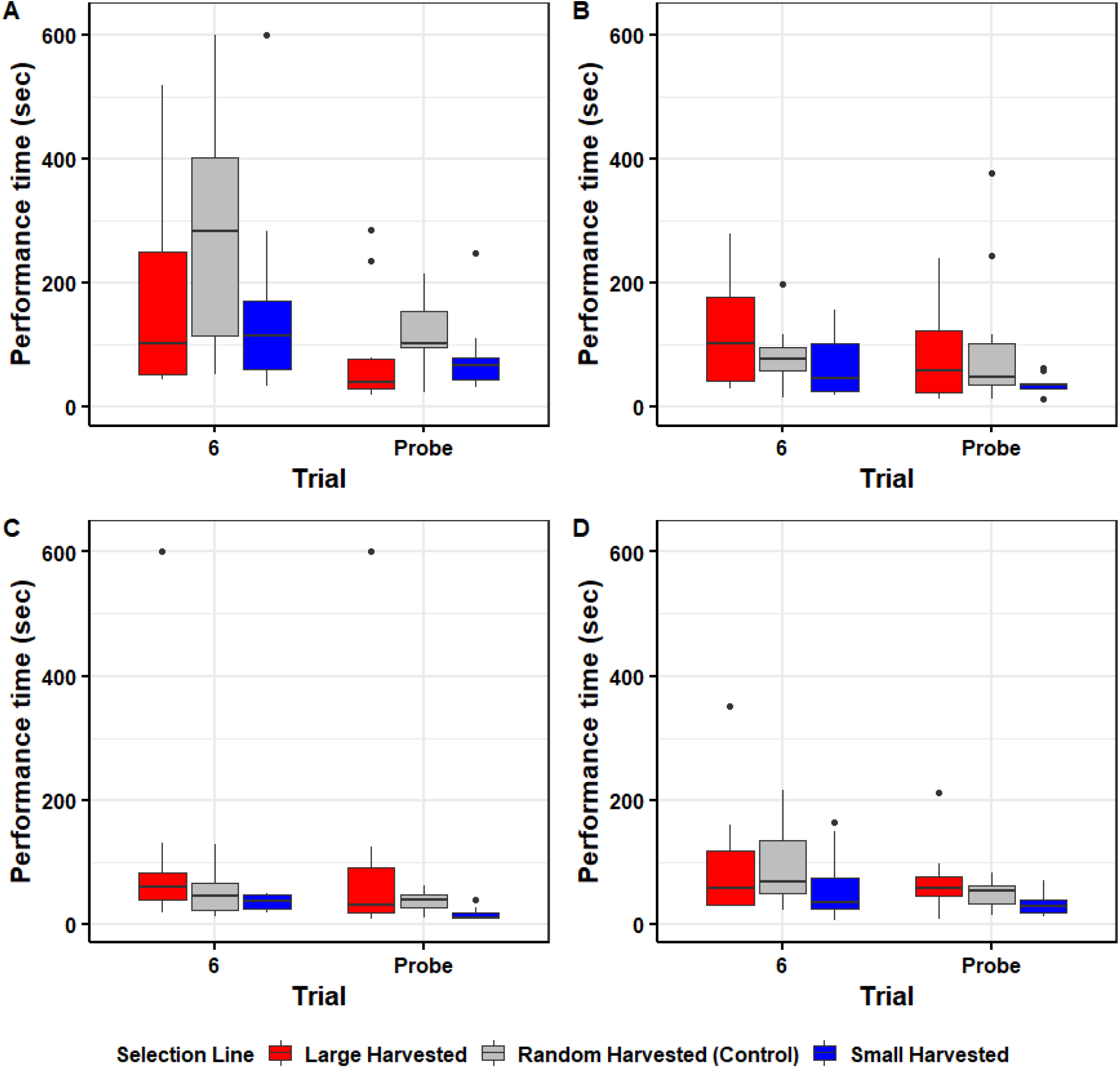
Comparison of performance time of fish across selection lines on 6^th^ and 12^th^-day (probe) trials as (**A**) juveniles (27-38 dpf), (**B**) subadults (69-80 dpf) and adults at (**C**) 112-123 dpf, and (**D**) 153-164 dpf

**Figure 4b:**
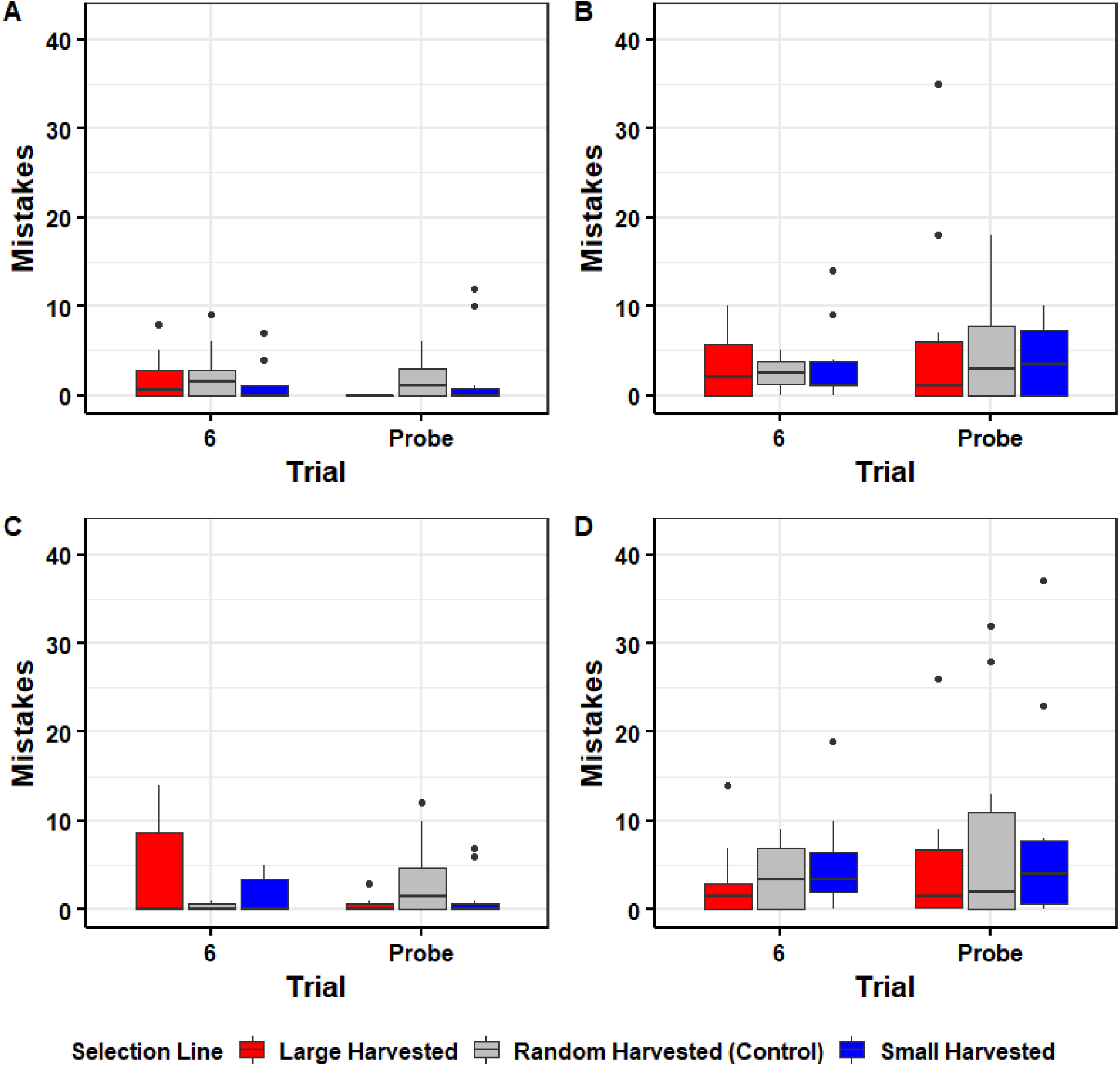
Comparison of mistakes made on 6^th^ and 12^th^-day trials as (**A**) juveniles (27-38 dpf), (**B**) subadults (69-80 dpf) and adults at (**C**) 112-123 dpf, and (**D**) 153-164 dpf

In the reversal learning assay, fish across selection lines showed no evidence of associative reversal ability (Fig. 5) because the percent time spent in rewarded and unrewarded chambers by fish were not significantly different (Table 4). In tests for collective decision making, fish generally made significantly faster decisions to enter the reward chamber over successive trials (F_1,147_=27.22, p<0.01; Fig. 6a) but we did not see significant differences across selection lines (F_2,147_=1.05, p=0.35). During the probe trial, we found that the large-harvested line fish took significantly less time to make decisions (by entering the correct door almost immediately after their release) compared to the 6^th^ trial (t=-3.54, p<0.01; Fig. 6b), while the small-harvested line fish did not differ in decision making time during 6^th^ and probe trials compared to the control line (t=-0.96, p=0.34; Fig. 6b). We summarize the results of all assays in Table 5.

**Table 4.** Paired t-test results for evaluating learning across selection lines during associative and reversal trials.

| Trials | Selection line | t | df | p-value |
| --- | --- | --- | --- | --- |
| Associative | Large-harvested | 1.54 | 8 | 0.16 |
|  | Random-harvested<br>(Control) | -1.19 | 8 | 0.27 |
|  | Small-harvested | -0.66 | 9 | 0.52 |
| Reversal | Large-harvested | 0.46 | 8 | 0.66 |
|  | Random-harvested<br>(Control) | -1.18 | 8 | 0.27 |
|  | Small-harvested | 0.34 | 9 | 0.74 |

**Table 5.**
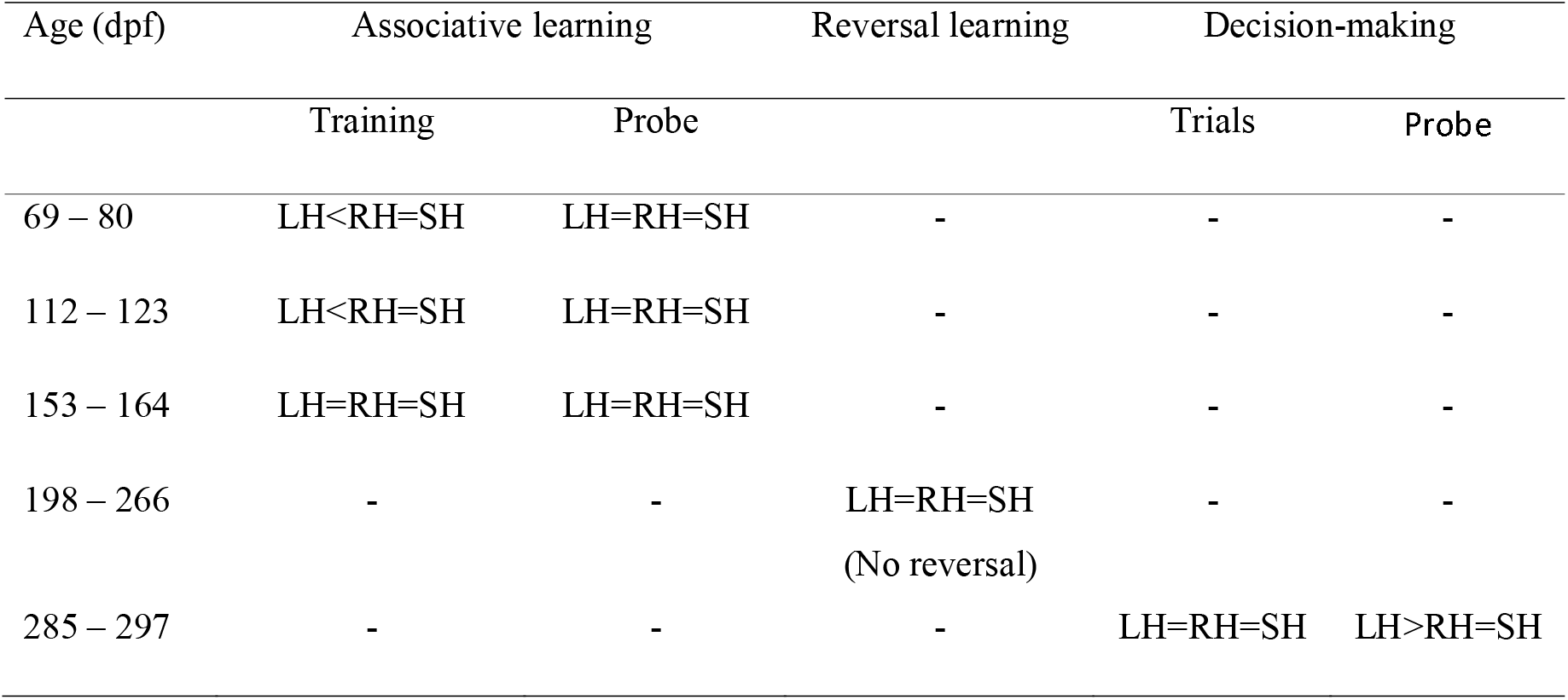
Summary of results of associative and reversal learning and collective decision-making assays. The symbols ‘>‘, ‘<’ or ‘=’ mean better, worse or similar performances.

**Figure 5:**
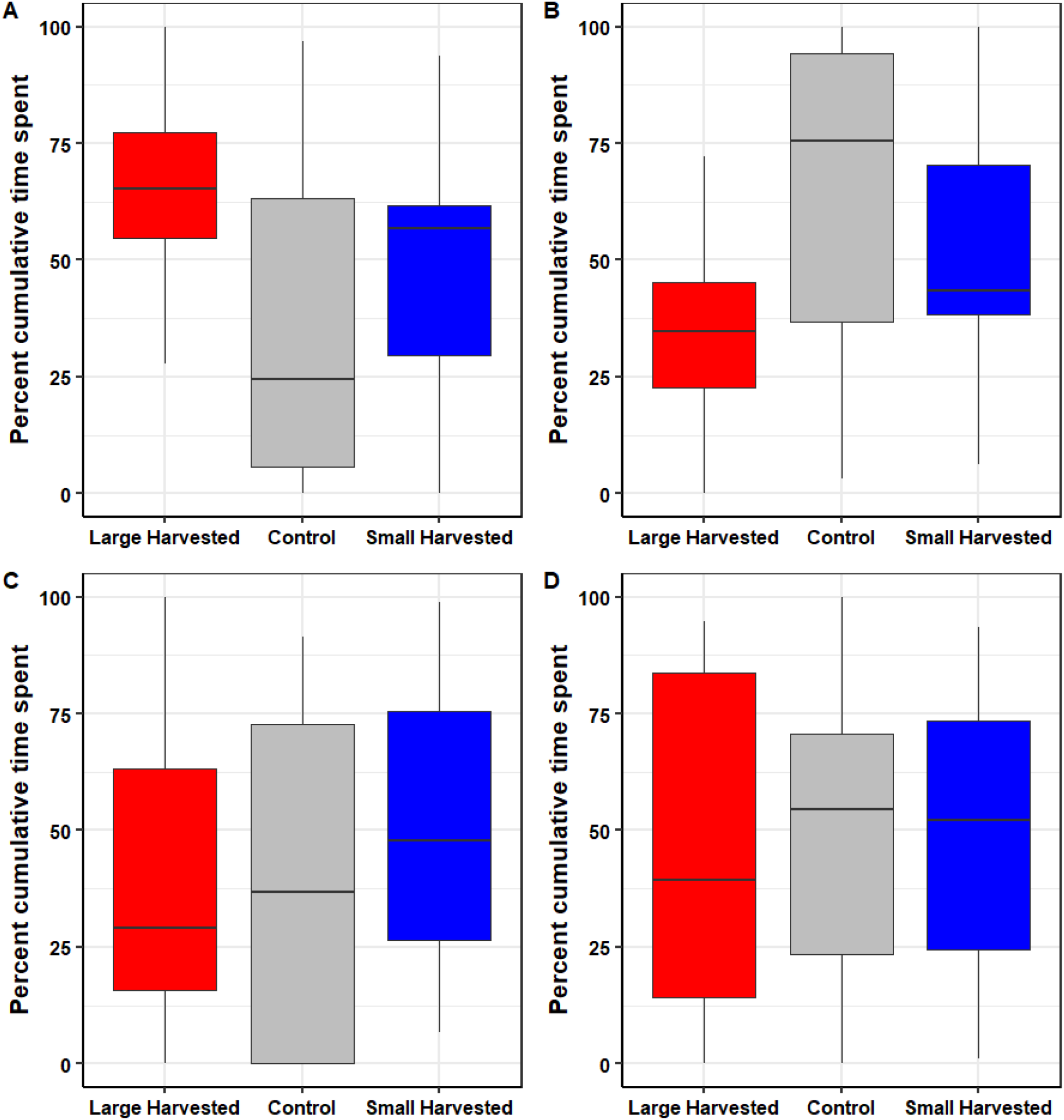
Results of reversal learning assay. Panels A and B show time spent in rewarded and unrewarded chambers during associative trials and panels C and D show time spent in rewarded and unrewarded chambers during reversal trials

**Figure 6:**
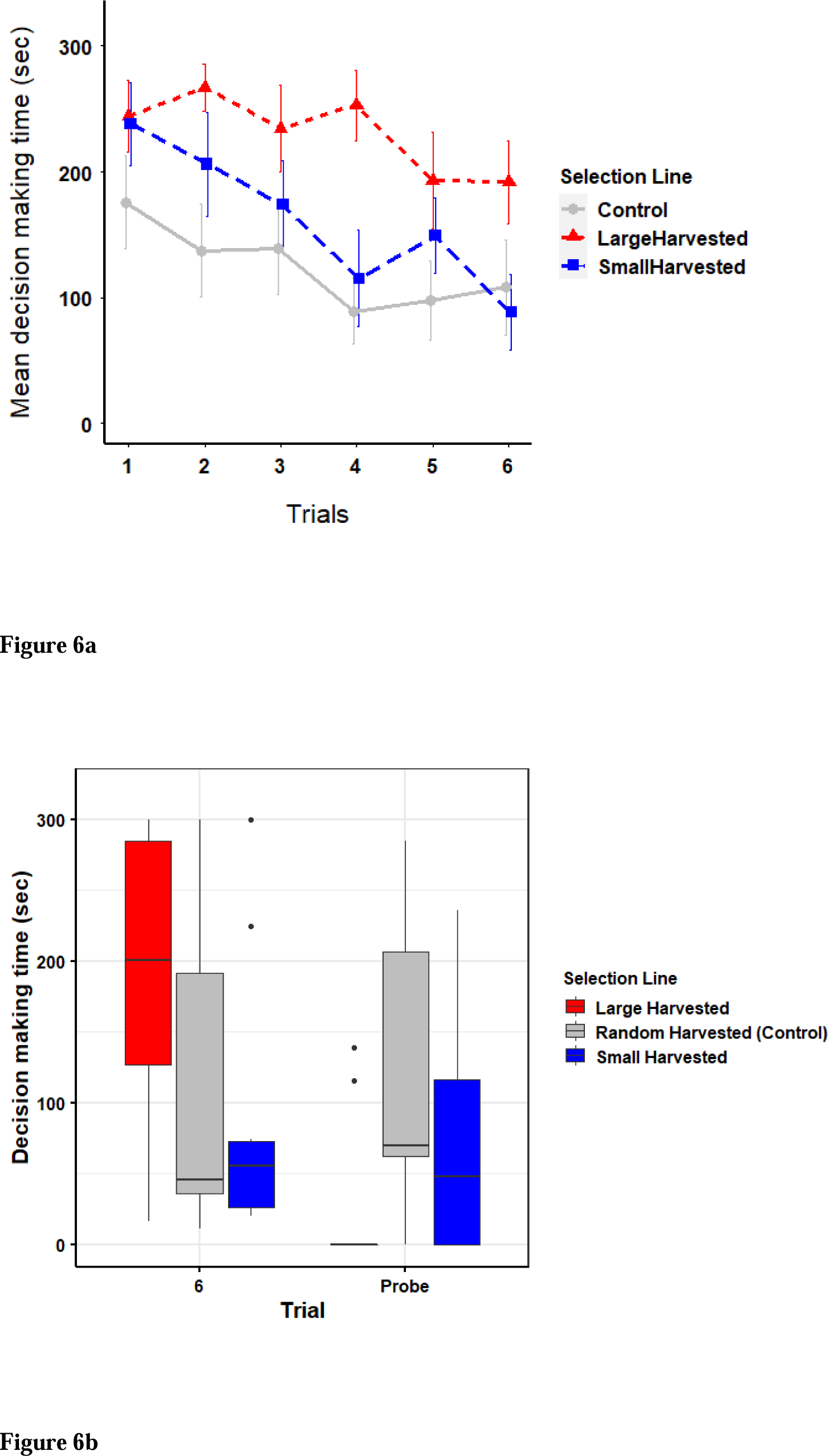
Comparison of (a) mean (±SE) decision-making time (sec) across trials, and (b) decision-making time during 6^th^ and 13^th^-day trials

## Discussion

Size-selective harvesting can have evolutionary consequences for adaptive behavioural traits (Andersen et al. 2018, Roy et al. 2021a, Sbragaglia et al. 2020). Here we provide evidence that size-selective harvesting can also evolutionarily alter learning and decision-making abilities in fish in ways that have adaptive value. We found that five generations of intensive size-selective harvesting followed by ten generations of no further selection had a substantial impact on associative learning and group decision-making in zebrafish. Performances of fish across selection lines improved significantly with ontogenetic age. Supporting our expectations, the positive size-selective harvesting resulted in slower learning speed in the subadults and adults compared to the controls. Yet, contrary to expectations, the selection lines did not show any reversal ability. Positively size-selected fish also collectively made speedy and accurate decisions when tested for associative memory. The slower learning speed shown by large-harvested lines can have relevant consequences for survival, but the improved collective decision-making ability might help the fish cope with certain threats. Further work is needed to test whether the selection lines differ in natural mortality rates against different predators.

We found that learning abilities (indicated by a decrease in the time to find a food reward and the number of errors) increased throughout ontogeny in all selection lines, as would be predicted (Spear et al. 2014). Older fish were significantly faster in locating the food reward even though we increased the size and/or complexity of the maze size at every ontogenetic time point. This could be due to two reasons. With the development of brain, fish are typically quicker in learning the location of the food reward (Spear et al. 2014). Alternately, swimming speeds of fish increase with ontogenetic age (Muller et al. 2000, Müller 2020) and fish could have reached the rewarded door faster as they grew in size. But this can only be a part of the explanation because the fish also made fewer mistakes across trials as adults, indicating better learning. Next, we found that the large-harvested line fish had slower learning speed as subadults and adults in a simple associative task. This agrees with our expectation that the large-harvested fish that exhibit a fast life-history would show lower learning ability due to possibly lower brain investment and shy behavioural tendencies. Our result can be explained based on the energy-tradeoff hypothesis (Isler et al. 2006, Isler et al. 2009), and may also be explained by the systematic relationship between learning ability and personality traits (Dougherty et al. 2018). Increased reproductive investment may be traded-off with decreased investment in expensive body-tissues like brain (Isler et al. 2006, Isler et al. 2009). Positive size-selection fosters the evolution of a fast life-history characterized by increased reproductive investment (Arlinghaus et al. 2009, Renneville et al. 2020, Uusi□Heikkilä et al. 2015) and this could have resulted in a decreased investment in brain development. Reduced brain investment may result in decreased learning abilities (Kotrschal et al. 2015, Kotrschal et al. 2013). Thus, the impact of size-selection on brain investment and neurological density is worthy of further investigation.

Personality traits like activity, exploration and boldness also relate to learning tendencies (Dougherty et al. 2018, Sih et al. 2012), and bolder, more exploratory and active animals were found to be faster in encountering and recognizing environmental contingencies and memorizing them (Gallistel et al. 2004, Griffin et al. 2015). The large-harvested line fish demonstrated shy behavioural tendencies as adults (Roy et al. 2021a, Sbragaglia et al. 2020) and this could explain why they showed slower learning abilities. By contrast, the small-harvested line fish were significantly bolder post the larval stages compared to the control and large-harvested line fish (Roy et al. 2021a, Sbragaglia et al. 2020). In our experiments, the small-harvested line fish were also more exploratory compared to the other two lines because they were quicker in finding the food reward on the first day of training (Trial 1, Fig. 3a) when the fish were exposed to the novel maze for the first time. Higher boldness and exploratory tendencies could explain why the small-harvested line fish took less time to find the food reward initially. However, higher boldness would mean that the fish might not thoroughly sample the arena and commit more errors in the process of doing so. This could be the reason why errors made across trials in the 4-month adults of the small-harvested line fish changed slowly even though they located the reward faster. We used a four-colour discrimination paradigm to test associative learning in adults at 154-164 dpf. The relatively higher errors during initial trials followed by a sharp decline in errors later in the small-harvested line fish could be because there were more chambers to explore.

We did not find evidence of reversal ability among selection lines. We used a social reward and tested reversal learning ability in groups of zebrafish. Because the fish were held and tested in groups, it was perhaps not motivating enough for the fish to locate the social reward. This is in contrast with a previous study where zebrafish successfully learned the location of a social reward and reversed their learning when the reward contingencies were changed but the fish were tested as individuals instead of groups (Roy et al. 2019). Further investigations with alternative rewarding paradigms could reveal differences in reversal abilities among selection lines. Roy et al. (2019) found reversal learning ability in wild, but not lab-bred populations of zebrafish. Wild populations evolve and develop under changing environmental conditions that favour cognitive flexibility (Tello-Ramos et al. 2019) while lab-reared populations develop under constant conditions. Here we used lab-bred selection lines of zebrafish that have been reared and maintained under consistent conditions and this could also be a reason why we did not find reversal ability among these fish.

We found that the fish across all selection lines made speedier decisions to enter the reward chamber over successive trials. This is similar to a pervious study in guppies where groups of fish made faster collective decisions to avoid a predator (Hansen et al. 2021). Improvement in decision-making speed over trials could be due to social facilitation (Brown et al. 2003) where one or more individuals enable others to find the food reward and the facilitation process speeds up over successive trials. The fish could also get more habituated with the setup over successive trials resulting in a decrease in decision-making time. The large-harvested line fish collectively entered the reward chamber significantly faster and almost instantaneously post-release during the probe trial compared to the 6^th^-day trial. This does not agree with our expectation that the large-harvested line will make slower decisions because they are less cohesive (Sbragaglia et al. 2021) and demonstrate higher within-group behavioural variability (Roy et al. 2021a). One explanation for this result could be that the social facilitation process was strengthened after the five-day interval period and the group members hastily followed the first individuals that entered the reward chamber. Another reason could be that fish copied choices made by others in the group instead of relying on personal information (Kendal et al. 2005) among the large-harvested line. Previous studies have shown that fish copy choices made by other members while foraging (zebrafish: Kadak et al. 2020, threespined sticklebacks *Gasterosteus aculeatus*: Pike et al. 2010). Because red is a preferred and attractive colour for zebrafish (Avdesh et al. 2012, Roy et al. 2019), one individual entering the red (reward) door might have led the other fish to copy this choice. This might have resulted in an instantaneous decision. Quorum responses by fish could have also increased the speed and accuracy among the large-harvested line (Ward et al. 2008). A threshold number of individuals could have chosen the more attractive stimulus and the rest of the fish would then have followed them leading to fast and accurate decisions (Sumpter et al. 2008, Ward et al. 2008).

In conclusion, our study provides experimental evidence for the evolutionary impact of size-selective harvesting on learning and decision making in fish. Positive size-selective harvesting that is typical of most commercial and recreational fisheries had the most significant impact on associative learning and group decision-making in zebrafish, reducing the learning speed and increasing the group decision making. These effects could affect growth and survival of exploited populations, because learning and making timely decisions are crucial for finding resources, avoiding dangers and adapting to changing environments. Further investigations are warranted to understand if the observed differences across selection lines also extend to other forms of learning and have a physiological underpinning, and whether they have survival consequences. Before such research becomes available, our work suggests that size-selective harvesting can evolutionarily affect the cognitive abilities of fish, which extends the evolutionary consequences of harvesting from life-history and behavior to the cognitive domain. One possible outcome could be an increase in natural mortality associated with reduced cognitive performance in fish exposed to positive size-selection, but whether this actually happens has to be examined in detailed predation experiments.

## Data Availability Statement

The data used in this study will be made available in the Dryad repository upon acceptance.

## Ethics statement

This study was approved by State Office for Health and Social Affairs Berlin (LaGeSo), Germany (approval number: G 0036/21).

## Conflict of interest statement

The authors declare no conflict of interest.

## Supplementary information

**Supplementary figure 1:**
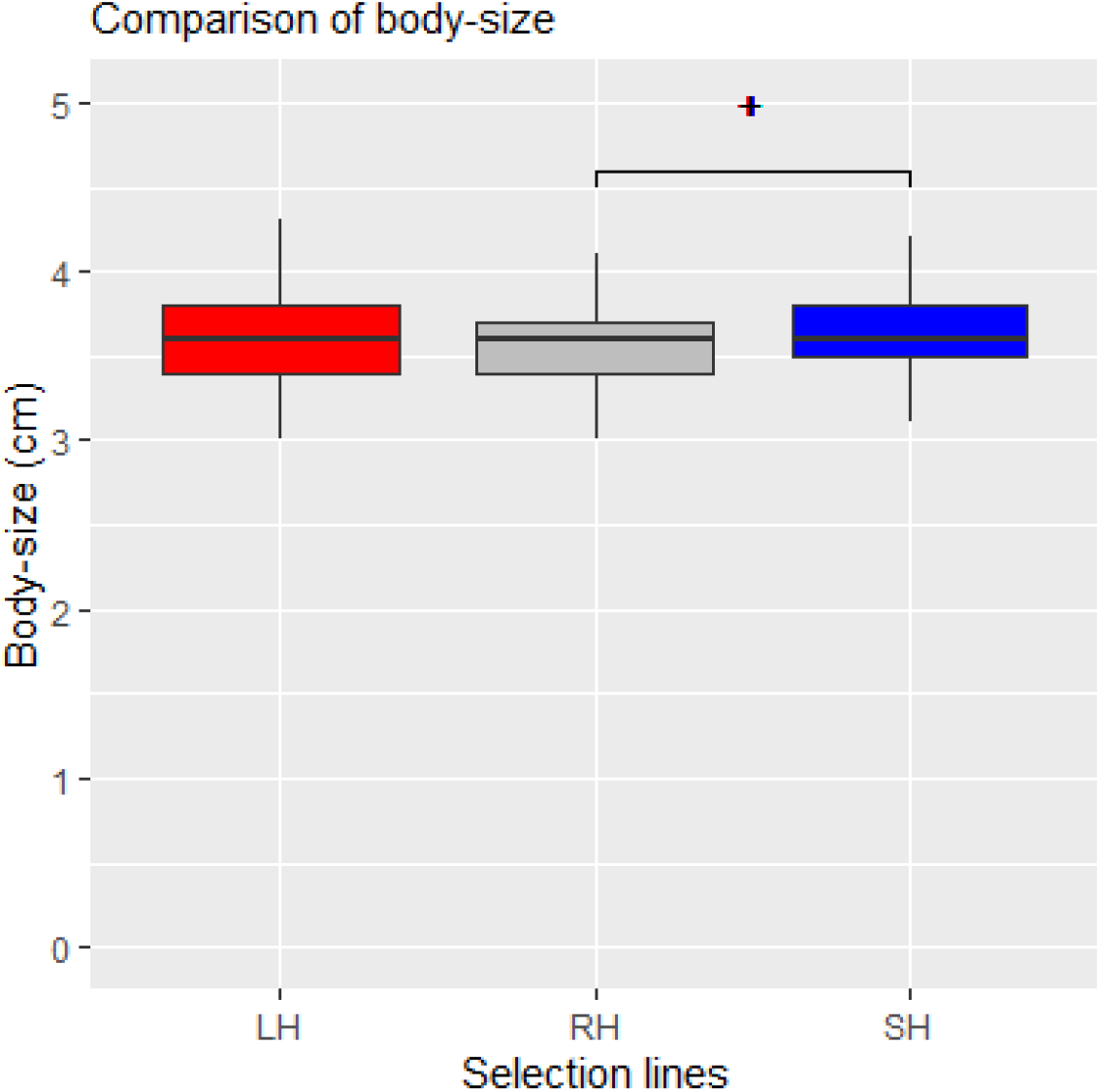
Comparison of body-size of non-experimental fish (11-month-old) across selection lines in F_16_. The small-harvested line (SH) fish were larger than the control (RH) line fish. Significant difference is indicated with code ^**+**^ (p=0.08).

## Notes

### Competing Interest Statement

The authors have declared no competing interest.

